# A One-Step Plasma Assisted Synthesis of Gold Nanoparticles and Simultaneous Linker-Free Conjugation with Nestin: An In Vitro Study of Cellular toxicity

**DOI:** 10.1101/2023.12.09.570950

**Authors:** Babak Shokri, Kimia Aalikhani, Melika Sanavandi, Mojtaba Shafiee, Hodjattallah Rabbani, Ghazaleh Fazli, Nilufar Sadeghi

## Abstract

We introduce a method for conjugating antigens to gold nanoparticles (GNPs) while synthesizing them using gas plasma, which eliminates the need for chemical linkers intended to facilitate the conjugation procedure for immunotherapy purposes. We report a physical approach to conjugate antigen Nestin (NES) as a marker in malignant tumors to GNPs. Two approaches were used to perform the conjugation of GNPs and NES. The first method involved using citrate to synthesize GNPs, and then NES was conjugated onto the GNPs surface by plasma. In the second method, GNPs were simultaneously synthesized and linker-freely conjugated to NES by plasma treatment. *Enzyme-linked immunosorbent assay* with the protocol defined in this study, *Zeta-sizer, Ultraviolet-visible spectroscopy*, and *Transmission Electron Microscopy* results confirmed NES conjugation to GNPs. In addition, the toxicity of the prepared samples was investigated in vitro using peripheral blood mononuclear cells (PBMCs) and *flow cytometry*, which proved the non-toxicity of the samples.

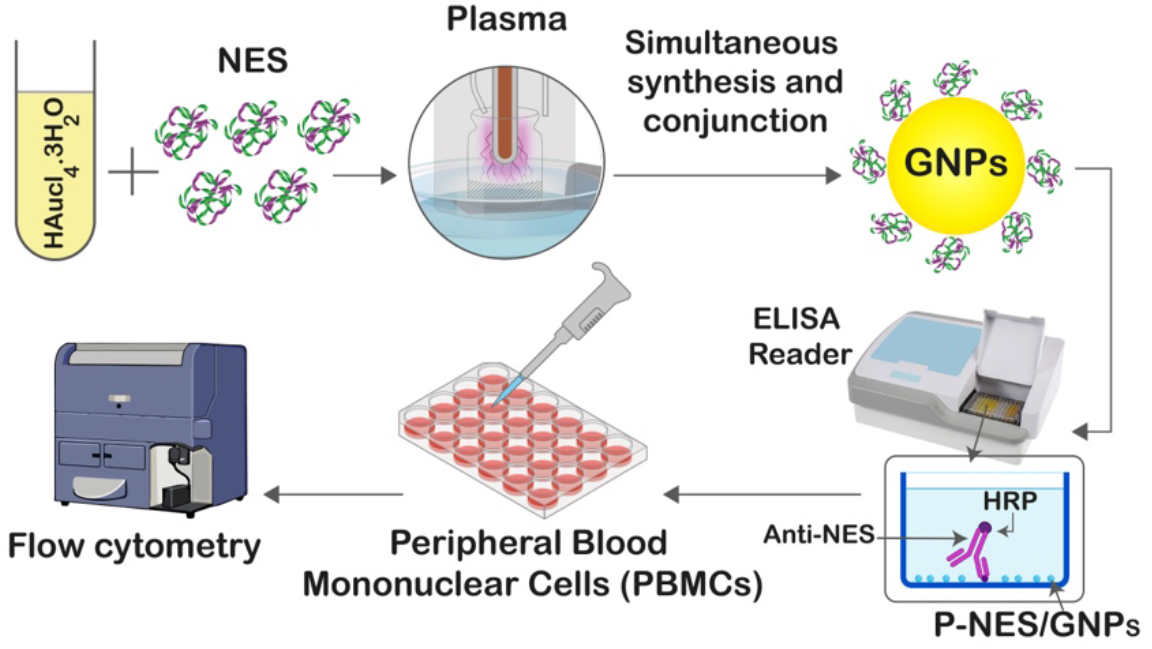

## INTRODUCTION AND BACKGROUND

Nanotechnology offers innovative strategies to regulate biomedical processes at the nanometer scale, poised to profoundly impact biology. Nanoparticles, particularly those with sizes comparable to molecules and biological structures, have garnered significant attention for their potential applications in biomedical research. [1] [2] Among the most biocompatible nanoparticles are GNPs, boasting various applications in modern medical and biological studies. [3]

GNPs are synthesized in three ways: biologically, chemically, and physically. Biosynthesis methods include the green synthesis of plant extract–GNPs and Green synthesis of Bacteria-GNPs [4]. The most well-known chemical methods are the citrate synthesis [5] and Frens Turkovich. [6] The newest physical methods are by laser [7] and gas plasma [8]. We used the citrate method and gas plasma in this study and compared the results at the end.

GNPs distinctive optical and physical properties [4], encompassing a large surface area to volume ratio [9], bio inertness [10], and high stability [11], make them valuable imaging agents in photoacoustic [12] and computed tomography (CT) imaging [13]. These properties, stemming from surface plasmon resonance [14] and bio-stability [15], further underscore their significance [16].

Extensive functionalization of GNPs with various biomolecules, including antibodies [17], peptides [18], and nucleic acids [19] has enabled targeted drug delivery to cancer cells, potentially minimizing side effects compared to traditional chemotherapy [20]. Notably, GNPs exhibit a high loading capacity with multiple ligands (chemical and biomolecules) [21]. In this study we used NES, a type VI intermediate filament protein [22] as a ligand which is mostly expressed in neural stem and progenitor cells during embryonic development and neurogenesis [23] Its versatile expression is notably evident in various tissues, including neural tissue [24], where it plays a pivotal role in the structural organization of neural precursor cells, contributing significantly to the development of the nervous system [25]. Beyond neural tissue, NES makes its presence known in blood vessels [26], specifically in the endothelial cells during embryonic development and under certain pathological conditions such as tumor angiogenesis [27] —signifying its involvement in the formation of blood vessels within tumors. [28] NES expression is often associated with more aggressive tumor behavior in cancer [29] and has been observed in a variety of cancer types including; brain tumors (often expressed in gliomas) [30], breast cancer [31], pancreatic cancer [32], prostate cancer [33], gastrointestinal cancers [34] (especially colorectal cancer) [29], and ovarian cancer

[35]. The presence of NES in various cancer types has prompted researchers to investigate its potential as a marker with implications for diagnosis and prognosis [36]. Notably, high levels of NES expression, detected through techniques like immunohistochemistry, may signify a more aggressive cancer phenotype, serving as a valuable diagnostic tool [37]. NES’s relevance extends to tumor grading and prognosis, where its elevated expression is associated with advanced and aggressive tumors [38]. This association provides critical information for predicting the likely course of the disease, guiding clinicians in determining appropriate treatment strategies. In the context of angiogenesis, NES’s expression in endothelial cells becomes crucial. As tumors necessitate a blood supply for growth, the detection of NES in tumor-associated blood vessels offers insights into the angiogenic potential of a tumor [39]. This multifaceted role of NES underscores its significance in unraveling the complexities of cancer biology and holds promise for enhancing diagnostic and prognostic approaches in oncology [36].

**Scheme 1.**
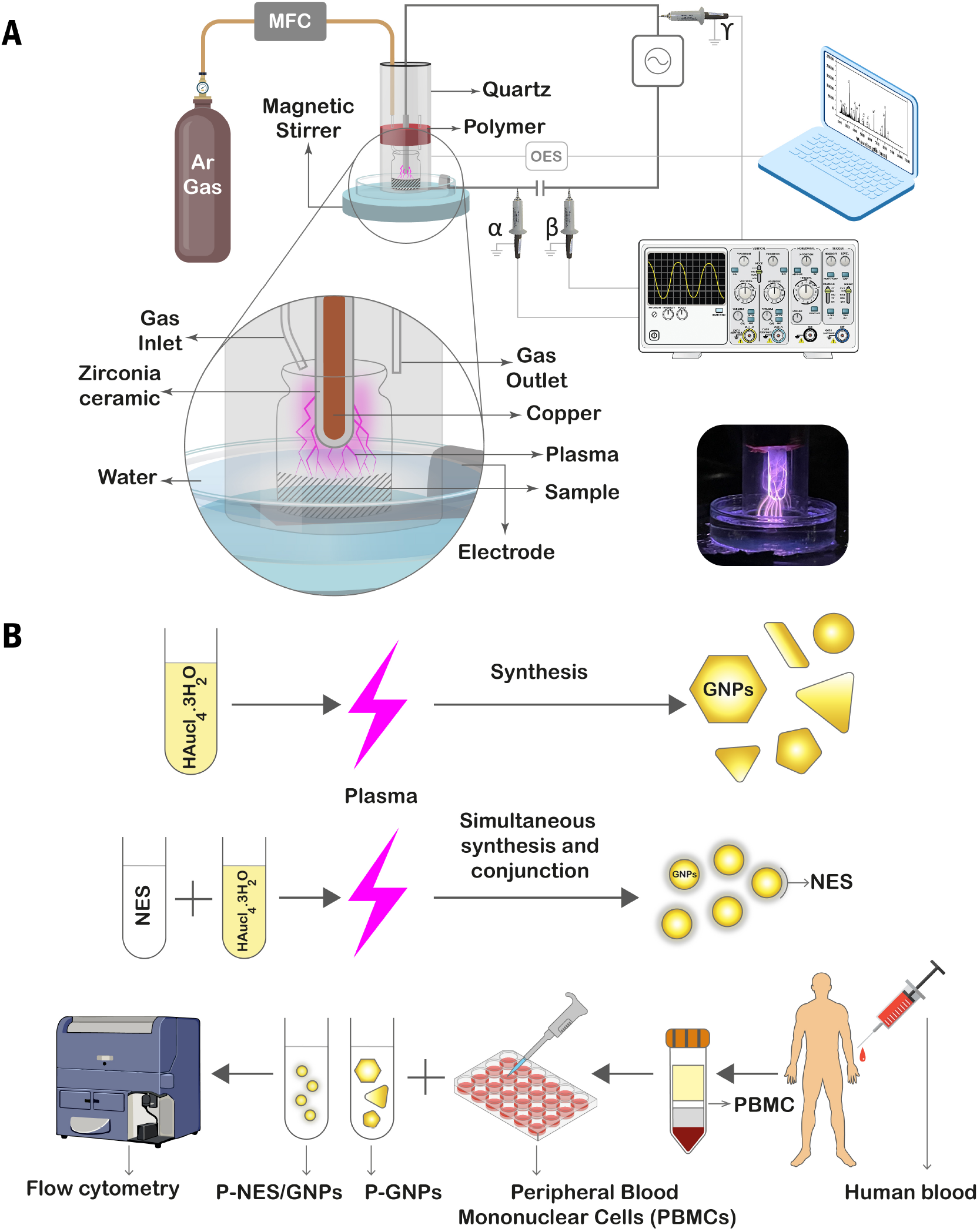
(A) The DBD plasma generator’s schematic with Argon gas consisting of a cylindrical quartz and a central electrode inside the cylindrical. The central electrode consists of a cooper nucleus, which is covered by zirconia ceramic. The oscilloscope high voltage probes were located in two positions: firstly, in α and β positions, then in β and γ to plot the Lissajous curve and calculate the power of plasma. The OES detector was located in two positions: near the electrode and the sample surface. (B) the schematic shows two pathways used in our study to synthesize plasma-assisted GNPs. Both groups were cultured with PBMC, which were isolated from human blood, to investigate the toxicity.

Bio-conjugation, which is an immobilization approach for connecting biomolecules to GNPs, can be achieved through physical or chemical methods. Physical interactions involve factors like hydrophobic or ionic attraction between GNPs and biomolecules, while chemical interactions employ methods such as chemical adsorption, bifunctional linkers, or adapter molecules like Streptavidin and Biotin. Linker chemistry is a common technique, though it has a lot of limitations and challenges [40].

A successful bioconjugation strategy requires controlled aqueous reactions [41] with stoichiometric efficiency, no disruptive agents, and selective bonding between linkers [42]. Linker stability is also important as it influences the release of the payload and modulates the toxicity [43]. Linkers should be easily incorporated into biomolecules, allowing simple quantitation [44]. The resulting bioconjugate should be stable across pH [45] and temperature ranges [46]. Challenges for Protein Bioconjugates include lack of site specificity, limited access to reactive sites, and difficulties with certain amino acids [47]. Additionally, chemical linkers may induce immunogenic side effects [48], necessitating careful consideration for biomedical applications. Numerous studies explore GNPs conjugation for various applications, addressing limitations to enhance effectiveness in cancer treatment and immunotherapy applications[49]. In a convergence of disciplines, in this study, we aimed to conjugate NES onto GNPs, addressing these challenges. The early part of this work involves the synthesis of citrate-stabilized GNPs. We utilized a protocol that enabled the preparation of approximately 10 nm – 100 nm spherical particles. The obtained particles were successively characterized using Zeta-sizer, Ultraviolet-visible spectroscopy (UV-Vis), and Transmission Electron Microscopy (TEM).

## RESULTS AND DISCUSSION

We performed a one-step linker-free plasma-assisted method to conjugate NES as a marker in malignant tumors to GNPs for immunization applications. We synthesized GNPs via two different methods: Plasma-assisted GNPs (P-GNPs) and Citrate-capped GNPs (C-GNPs).

The primary gold solution for these two groups was prepared by Tetrachloroauric (III) acid ( HauCl_4_. 3H_2_O ) with 40 mg/ml concentration purchased from SHIRAZ CHEM. Afterwards, we prepared a solution of 125 μl HauCl_4_. 3H_2_O then we brought the solution to 50 ml by adding distilled water and we used this to prepare our experiment samples. C-GNPs with size around 30 nm were prepared based on the previous study protocol. (Kumar, Aaron et al. 2008) Trisodium citrate was purchased from MERCK. In the following, we will explain the protocol we used to prepare P-GNPs:

To synthesize P-GNPs, we designed three different atmospheric plasma generators (see details in supporting information, Section 1). Eventually, after considering the temperature condition, we chose the dielectric-barrier discharge (DBD) plasma set-up constructed at the Laser and Plasma Research Institute (LAPRI) at Shahid Beheshti University. The device used in this study consisted of two main parts, namely the electrode and high voltage power supply. The electrode was constructed using zirconia ceramic with a copper core measuring 5 cm in length and 7 mm in diameter. We used argon gas with a 1Lpm flow and a pulse mode source to produce plasma (Scheme 1). The timing of plasma treatment was optimized to 15 minutes as an appropriate and efficient time to synthesize P-GNPs. We prepared a solution of 1.5 ml from our primary gold solution and 1.5 ml of distilled water; then, we treated the solution via plasma to synthesize P-GNP.

All the plasma-treated ingredients were exposed to plasma for 15 minutes according to the optimal time. The distance between all samples and the zirconia-covered electrode was 1 cm.

We determined the power of plasma by the Lissajous curve (Figure 1A). [50] We engineered the circuit to obtain a stable condition for temperature control. Since Temperatures up to 50° C do not damage the protein structure, we designed the setup to keep the temperature below 50° C (Figure 1B). [51] Additionally, we assayed the molecular weight of NES after plasma treatment by SDS-page analysis, which showed no difference (Figure 1C).

**Figure 1.**
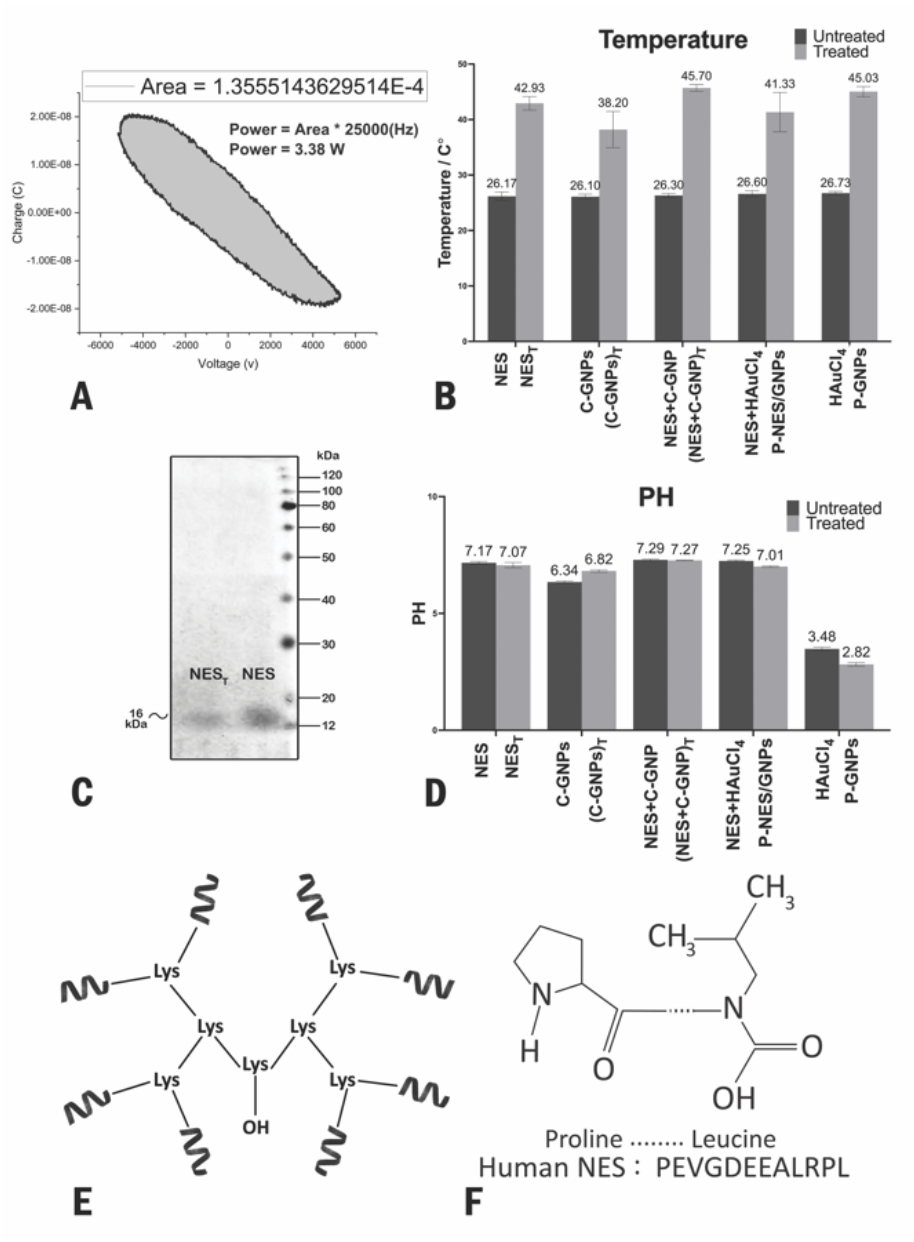
(A) Lissajous curve, which shows the power of plasma. (B) Temperature changes chart for NES, C-GNPs, NES+C-GNPs, NES+ HauCl_4_ and HauCl_4_after 15 min plasma treatment. (C) SDS-page analysis for NES and plasma-treated NES indicates no molecular weight difference (∼16 kDa) occurred by plasma. (D) PH changes chart for NES, C-GNPs, NES+C-GNPs, NES+ HauCl_4_and HauCl_4_ after 15 min plasma treatment. (E) Structure of NES (8MAPs). (F) NES sequence.

The reactive species generated by the plasma were characterized using optical emission spectroscopy (OES) at two different positions, close to the electrode and near the sample surface (Figure 2C).

**Figure 2.**
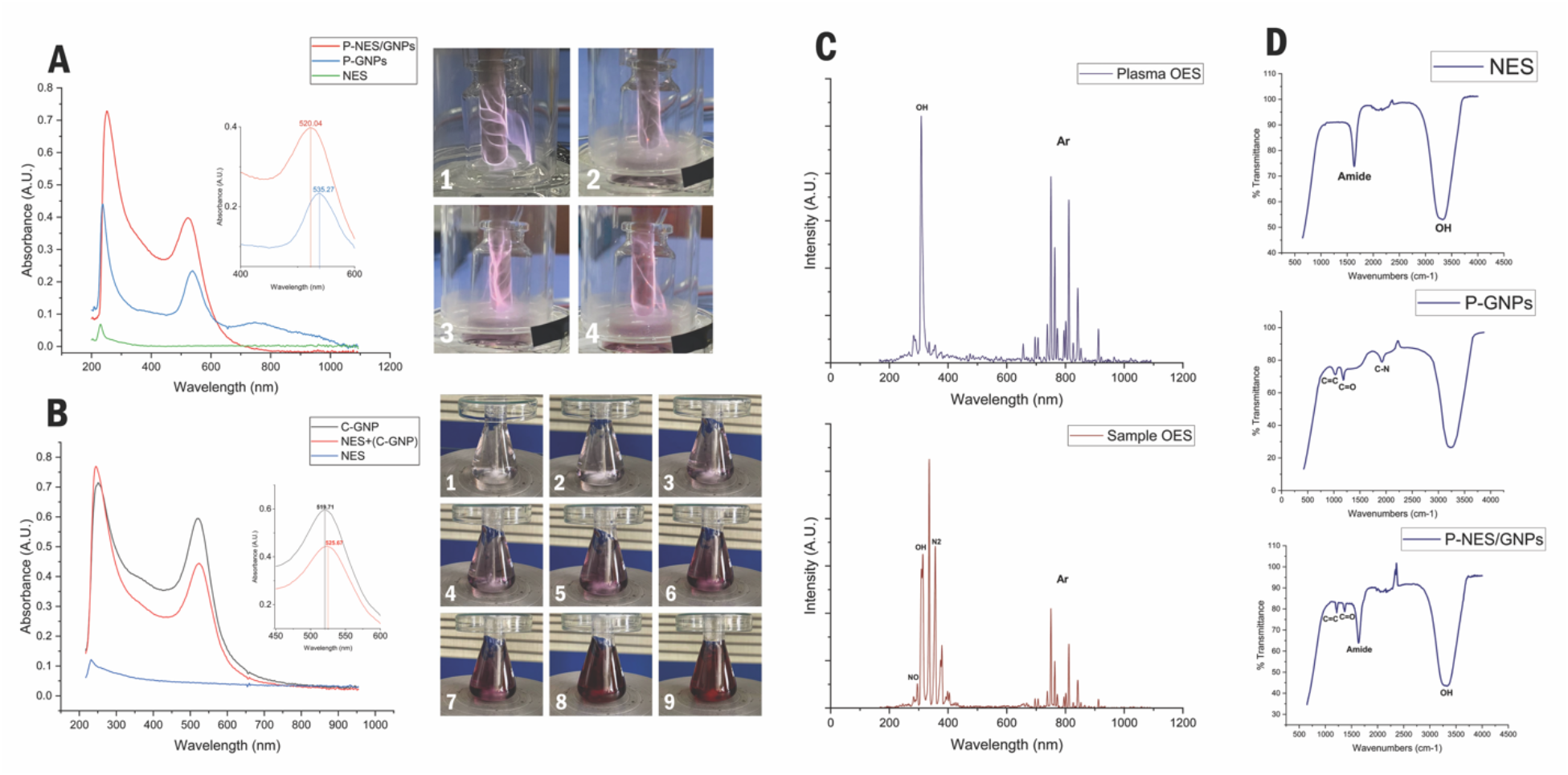
(A) UV-vis of P-NES/GNPs, P-GNPs and NES. The simultaneous conjugated and synthesized P-NES/GNPs shows an absorbance wavelength peak of 520.04 nm, which indicates particle size is around 25-30 nm. However, the P-GNP peak stands at 535.27 nm, indicating that the size of particles is around 100 nm. The pictures 1-4 show changes in solution color during plasma treatment. (B) UV-vis of NES +C-GNPs, C-GNPs and NES. NES +C-GNPs has an absorbance peak in 525.67 nm wavelength. C-GNPs peak stands at 519.71 nm. Here, we used the same GNPs, so the shift observed in UV-vis (which indicates the size change) confirms the conjugation of NES to C-GNPs. The pictures on the right 1-9 show the color change steps of C-GNPs during synthesis. (C) The result of the OES analysis shows that the concentration of Ar near the electrode is higher, and there is an OH peak near the sample because of evaporation. (D) the result of FTIR-ATR shows functional groups of NES, P-GNPs and P-NES/GNPs. We see the presence of carbonyl group in P-GNPs which is eliminated in P-NES/GNPs which has given its place to Amide Group, confirming the successful conjugation along with other tests results.

To prepare our experiment samples to investigate the linker-free conjugation, we proceeded as follows. Synthetic Nestin (NES) 8MAPs peptide was ordered from PADZA company with 100 μg/ml concentrations (in PBS solution) [52], which subsequently will be mentioned as the primary NES solution to conjugate it on our nanoparticles. NES molecular weight (∼16 kDa), schematic structure, and sequence are shown in Figures 1C, E, and F, respectively.

Samples were prepared in 3 ml with three repetitions on three days.

We defined our experiment groups below, which are also shown in Table 1:

**Table 1.** Experimental samples.

|  |  |
| --- | --- |
| Group 1 | NES + C – GNPs |
| Group 2 | NES<sub>T</sub> + C – GNPs |
| Group 3 | NES + (C – GNPs)<sub>T</sub> |
| Group 4 | (NES + C – GNPs)<sub>T</sub> |
| Group 5 | P – NES/GNPs |
| Group 6 | NES + P – GNPs |
| Group 7 | NES<sub>T</sub> |
| Group 8 | NES |
| Group 9 | (C – GNPs)<sub>T</sub> |
| Group 10 | P – GNPs |
| Group 11 | C – GNPs |

- Group 1: 1.5 ml of primary NES solution was added to 1.5 ml C-GNPs with no plasma treatment to investigate the spontaneous conjugation.
- Group 2: At first, 1.5 ml of primary NES solution was treated with plasma (to see if plasma treatment can activate the antigen and do the conjugation process [53]), and then it was added to 1.5 ml C-GNPs.
- Group 3: We added 1.5 ml C-GNPs, which was treated by plasma to 1.5 ml primary NES solution (to see if plasma treatment can activate the GNP surface and make the conjugation process easier).
- Group 4: We added 1.5 ml of primary NES solution to 1.5 ml of C-GNPs, and then we treated the whole solution via plasma to investigate the conjugation process.
- Group 5: In this group, we simultaneously synthesized GNPs and conjugated NES to them. We prepared our solution by adding 1.5 ml of primary NES to 1.5 ml of primary gold and then treated the final solution via plasma.
- Group 6: First, we synthesized P-GNPs by treating the primary gold solution, and then we added 1.5 ml of P-GNPs to 1.5 ml of primary NES solution to check their spontaneous conjugation.
- In this group, 1.5 ml of primary NES solution Group 7: was added to 1.5 ml distilled water, and then the final solution was treated with plasma to see if plasma treatment could change the size of our peptide.
- Group 8: We added 1.5 ml of primary NES solution to 1.5 ml of distilled water to have the same concentration as a positive control group.
- Group 9: First, 1.5 ml of C-GNPs was added to 1.5 ml of distilled water, then treated by plasma as a control group.
- Group 10: 1.5 ml of primary gold solution was added to 1.5 ml of distilled water, and then the whole solution was treated with plasma.
- Group 11: 1.5 ml of C-GNPs was added to 1.5 ml of distilled water as a control group.

All samples treated with plasma were 3 ml, and the evaporation during plasma treatment was compensated by adding the distilled water until the first volume was reached.

PH and temperature were measured for all plasma-treated samples before and after treatment. (Figure 1D) High-Performance Liquid Chromatography (HPLC) was also done to trace HCL, the results of which show that HCL is not present in samples as a plasma by-product, and therefore it has no effect on PH.

To survey the conjugation, each set of 11 samples was filtered thrice on three days using centrifugal filter units 30 KDa Mwco. By optimizing the centrifuge time to 20 minutes and 1000 RPM, we made sure most of the naïve NES passed the filter (see details in supporting information, Section 2). All the samples were 3 ml, and after filtration, the up and down of the filtered samples were reached to the volume of 3 ml by adding the background solvent. In the next step, ELISA was taken a week after from filtered (up and down) and unfiltered samples three times on three different days (Figure 3A and B). ELISA was conducted with a direct method, which involved coating our prepared samples as the first layer, and then HRP-NES antibody was added to ELISA wells to indicate antigen-antibody interaction (see details in supporting information, Section 3).

**Figure 3.**
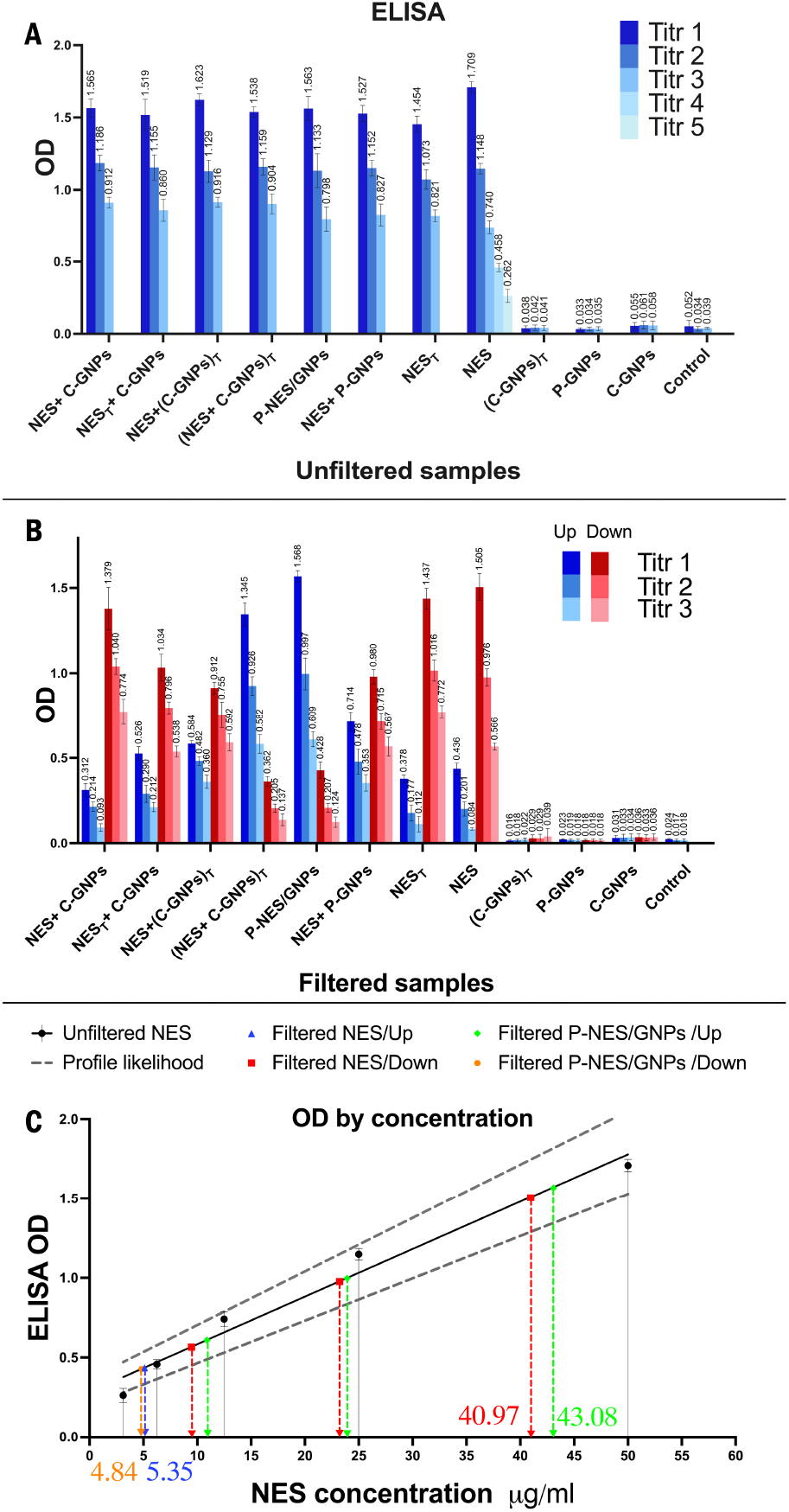
(A) ELISA results for all thrice 11 samples without filtration. All NES-included samples indicate nearly identical optical densities (ODs). For the naïve NES sample, we performed the assay in 5 titrations. Since the ELISA has been conducted with the direct method, the control group included BSA and anti-NES-HRP. (B) ELISA results for all thrice 11 filtered samples for both the up and down sides of the filter. (C) By using the OD of 5 titrations of unfiltered naïve NES, we plotted the OD by NES concentration graph to compare the up and down sides of filtered samples. This analysis revealed that the NES concentration of filtered P-NES/GNPs/Up is approximately 43 μg/ml.

Due to the analysis of ELISA results (Figure 3B), we demonstrated that the linker-free conjugation of group 4 and group 5 is successfully done. The ELISA test was repeated after two weeks and one month, and the same results were observed (see details in supporting information, Section 4).

The comparison of the OD of these two groups (4 and 5) shows us that the simultaneous synthesis and conjugation for the P-NES/GNPs group has the most OD (upside of the filter), which is the best result among other groups.

In unfiltered samples ELISA, we did the assay for naïve NES in 5 titrations, 50, 25, 12.5, 6.25, and 3.125 μg/ml, to plot the best-fit line in the “OD by concentration” graph (Figure 3C). This graph demonstrates the correlation between OD and NES concentration. The NES concentration for P-NES/GNPs and NES (up and down) are shown in Figure 3C. As we can see, the NES concentration of P-NES/GNPs (up) is approximately 43 μg/ml, whereas around 40 μg/ml of naïve NES has passed through the filter (see details in supporting information, Section 5).

The UV-vis results exhibit a peak shift in groups 4 and 11 (before and after conjugation), which can be equivalent to a change in particle size which is related to the presence of the protein (Figure 2B). [54] This shift may also arise from alterations in the dielectric coefficient [55], linked to the presence of the protein as well. Whether caused by changes in particle size or dielectric coefficient, this shift serves as confirmation of the successful conjugation. Furthermore, there is a significant peak shift which is observed for groups 5 and 10 (Figure 2A). The wide peak for group 10 in the region of 600 to 800 nm (Figure 2A) indicates the presence of GNPs in different forms. [56]

We had two different approaches for synthesizing P-GNPs. We synthesized them with and without NES. The TEM results (Figure 4A) confirm the nanoparticle shape variation predicted by UV-vis. The shapes of P-GNPs are not homogenous; the average size is ∼100 nm. Conversely, P-NES/GNPs are desirably homogenous and spherical, and the average size is ∼25-30 nm. Furthermore, there is a noticeable halo around P-NES/GNPs in TEM images (Figure 4A-4), which, based on previous studies [57], is another confirmation of successful conjugation and verified the UV-vis results.

**Figure 4.**
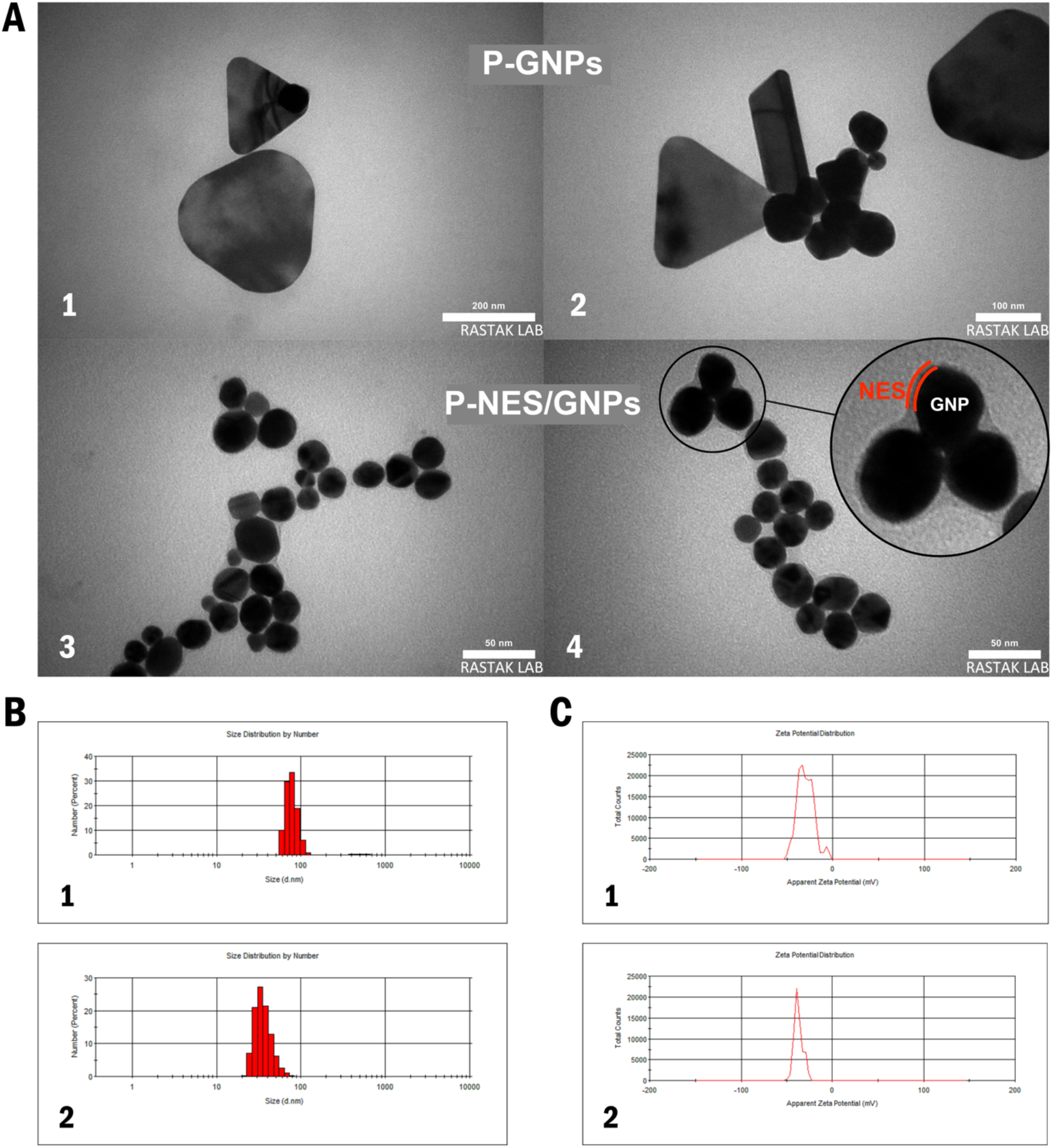
(A) Transmission Electron Microscopy (TEM) images of P-GNPs and P-NES/GNPs. Pictures 1 and 2 show the plasma-assisted synthesized GNPs, which exhibit amorphous shapes. Their average size is approximately 100 nm. Pictures 3 and 4 represent the plasma-assisted synthesized NES/GNPs. The P-NES/GNPs appear homogeneous and display a halo surrounding the particles. This halo in picture 4 evidences the conjugated NES. The average size of P-NES/GNPs is around 25-30 nm. (B) Zeta sizer results of size distribution by number. Pictures 1 depicts the size distribution by number of P-GNPs, which peaks at approximately 80 nm. Pictures 2 portrays the size distribution by number of P-NES/GNPs, exhibiting a peak between 20 and 30 nm. (C) The graphs represent the zeta potential distribution of P-GNPs and P-NES/GNPs. Graph 1 illustrates the surface charge of P-GNPs, while graph 2 shows that of P-NES/GNPs, demonstrating an increased negative surface charge.

The FTIR-ATR analysis of the GNPs group reveals the presence of the carbonyl group (Figure 2D). Conversely, in the case of the naive NES, the FTIR-ATR spectrum exhibits the amide group indicative of its proteinaceous structure. Following the FTIR-ATR of P-NES/GNPs, it is observed that the carbonyl group is eliminated, but the amide peak is still present along with other GNPs peaks. This test outcome, along with other tests, confirms the successful conjugation.

In Figure 4B and C Zeta-sizer results are shown. We can see size distribution by number results for P-GNPs and P-NES/GNPs in Figure 4B-1 and 2 respectively. Which indicates the particles size peak for P-GNPs around 80-100 nm and the peak for P-NES/GNPs approximately in 20-30 nm. The zeta potential test results (Figure 4C-1,2) indicate an increased level of negative charge in the conjugated group compared to bare nanoparticle, signifying the presence of the protein (NES) on nanoparticle’s surface. Bare GNPs had negative charge themselves and from these results we can determine that the positive functional group of NES is attached to the nanoparticle, and its negative functional group remains free (see details in supporting information, Section 6).

In the final step, we assessed the toxicity of our prepared samples, including P-NES/GNPs, naïve NES, treated NES and P-GNPs. Peripheral blood mononuclear cells (PBMC) were isolated from three human blood samples for Annexin V flow cytometry. This test is a robust assay to determine the toxicity against immune blood cells. Human blood samples were centrifuged twice for 10 minutes and washed to isolate the PBMCs. Subsequently, the prepared PBMC was cultured (500’000 cells per well) with samples from groups 5, 7, 8, and 10 (95 μl each sample added to 5 μl PBS 20X). The apoptosis kit was purchased from PADZA PADTAN PAJHOOH, and control groups included auto-apoptosis, only PI, only FITC, and the unstained group. The results are shown in (Figure 5A and B). The obtained data may assist in determining the amount of NES to be used for in vivo immunization in pre-clinical studies.

**Figure 5.**
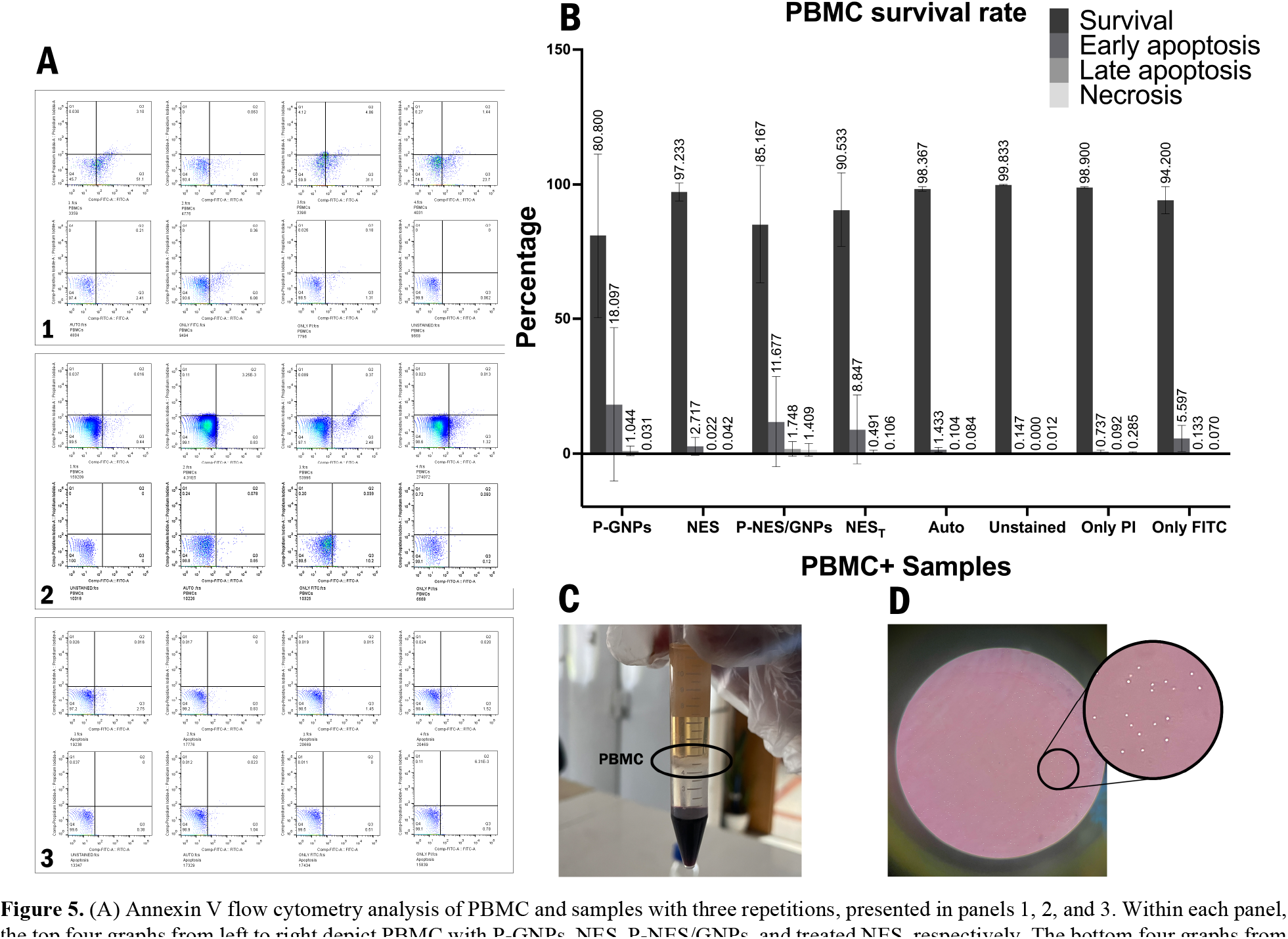
(A) Annexin V flow cytometry analysis of PBMC and samples with three repetitions, presented in panels 1, 2, and 3. Within each panel, the top four graphs from left to right depict PBMC with P-GNPs, NES, P-NES/GNPs, and treated NES, respectively. The bottom four graphs from left to right represent the control groups, including auto-apoptosis, unstained cells, only PI, and only FITC, respectively. (B) The graph compares the Annexin V flow cytometry results. Considering the number of cultured cells per well (500,000 cells), the survival rate of PBMC+P-NES/GNPs can be considered non-toxic. (C) The human blood sample after centrifugation, highlighting the PBMCs. (D) The microscopic image of a well containing PBMCs.

## CONCLUSION

We provided a facile, one-step method to synthesize GNPs and simultaneously conjugate them to NES antigen without using any chemical linkers, making the conjugation process significantly fast and cost-effective. We used NES as a stabilizer, which can be a tumor marker. Controlling the shape, size and stability of P-GNPs is a challenge. We demonstrate that using peptides, specifically antigen, helps synthesize homogenous GNPs and also has acceptable stability after months; furthermore, based on toxicity results, it has the immunization application potential.

As shown in this research, we propose that plasma can be a revolution in the field of nanoparticle peptide conjugation since it is a fast, linker-free, stable and cost-effective method.

To further explore this conjugation technique, applying it to other nanoparticles would be worthwhile. Our particular intention is to optimize the performance of GNPs in immunization, as this would represent a significant step forward in developing cancer treatments. In the next stage of our research, the most favorable samples based on ELISA results can be used to conduct in-vivo experiments for studying the performance, biocompatibility of samples and the immune system response.

## ASSOCIATED CONTENT

Supporting Information

The Supporting Information is available free of charge at:

Details of various plasma generators, filtration optimizing, samples stability, conjugation efficiency and Zeta-sizer results.

## Supporting information

see details in supporting information

## AUTHOR INFORMATION

## Author Contributions

Kimia Aalikhani and Melika Sanavandi contributed equally to this work as first authors.

### Notes

The authors declare no competing financial interest.

## ACKNOWLEDGMENTS

This work was kindly funded by PADZA PADTAN PAJHOOH.

