## Supplementary material for "A One-Step Plasma Assisted Synthesis of Gold Nanoparticles and Simultaneous Linker-Free Conjugation with Nestin: An In Vitro Study of Cellular toxicity": see details in supporting information

# of

**Section 1:**

**Plasma generators**

Our objective in this study was to develop a linker-free method for conjugating peptides to gold nanoparticles (GNPs) using gas plasma and make the conjugation process time and cost-effective. Initially, four different antigens, PeptideV1, RECV1YP, AOD-R16, all synthetic peptides from ROR1 antigen, and Nestin (NES), were used to be conjugated to GNPs.

We ordered all antigens from PADZA company with 100 µg/ml concentrations (in PBS solution); PeptideV1 was a monomer peptide of ROR1 antigen, RECV1YP was an 8MAPs peptide of ROR1 antigen, and AOD-R16 a 16MAPs peptide of ROR1. Ultimately, NES was selected due to its established role as a tumor marker and its promising therapeutic applications. In pursuit of optimized conjugation and GNPs synthesis, we used three distinct plasma setups: plasma jet, micro-discharge, and dielectric-barrier discharge (DBD). The schematic of jet and micro-discharge setups are given below, and the DBD setup schematic is already shown in the main text.

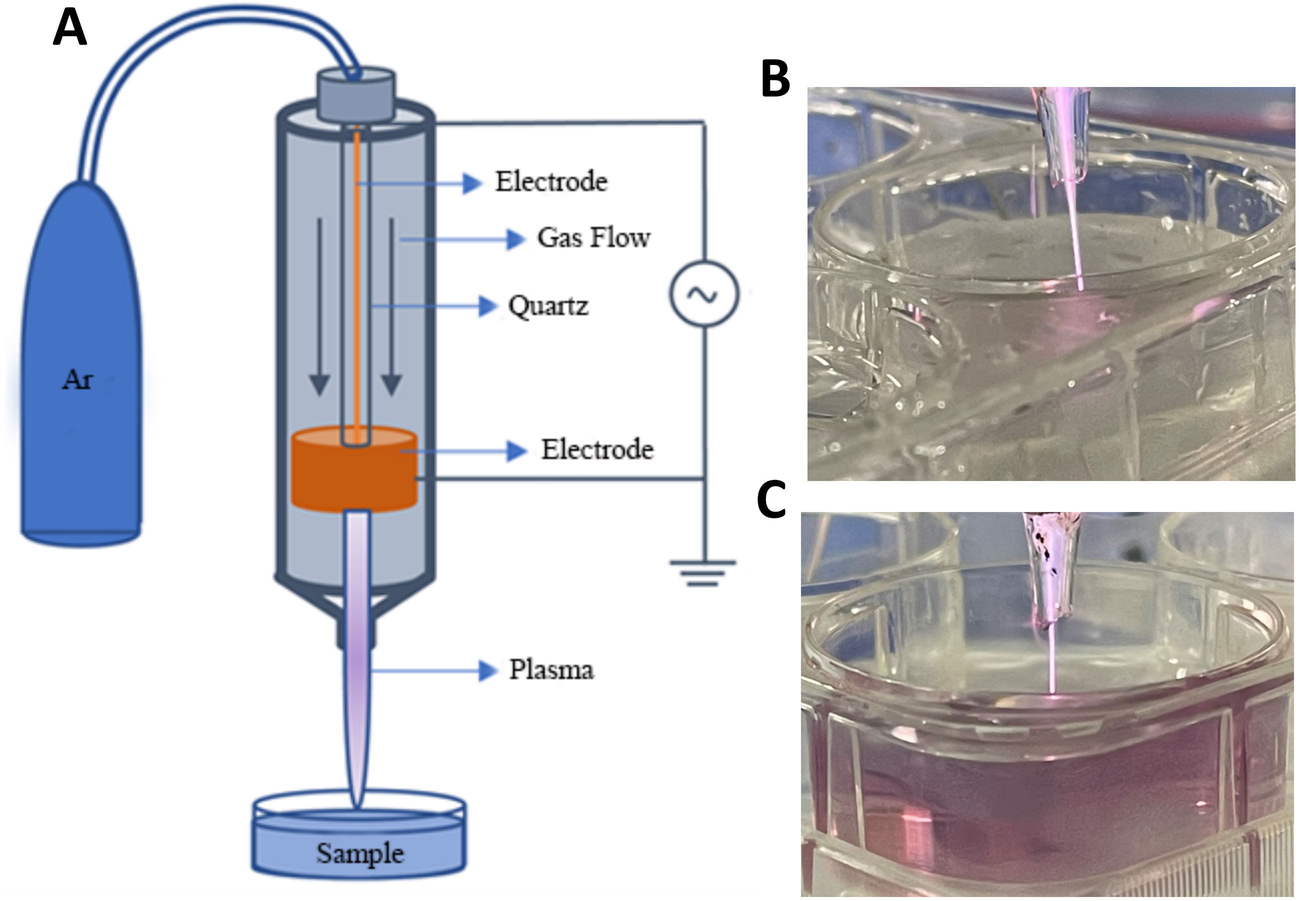


***Figure S1:*** *A) The plasma jet setup consists of a 100-millimeter-long quartz tube with an inner diameter of 5 millimeters and an outer diameter of 7 millimeters. Two copper tape electrodes, one powered (high voltage) and one grounded, which are wrapped around the external surface of the tube with a 10-millimeter gap. The electrodes are spaced 6 millimeters from the tube's exit nozzle. B) Treating the primary gold solution to synthesize GNPs. C) Here, we can see the color change of the solution, which indicates the synthesis of GNPs after plasma treatment.*

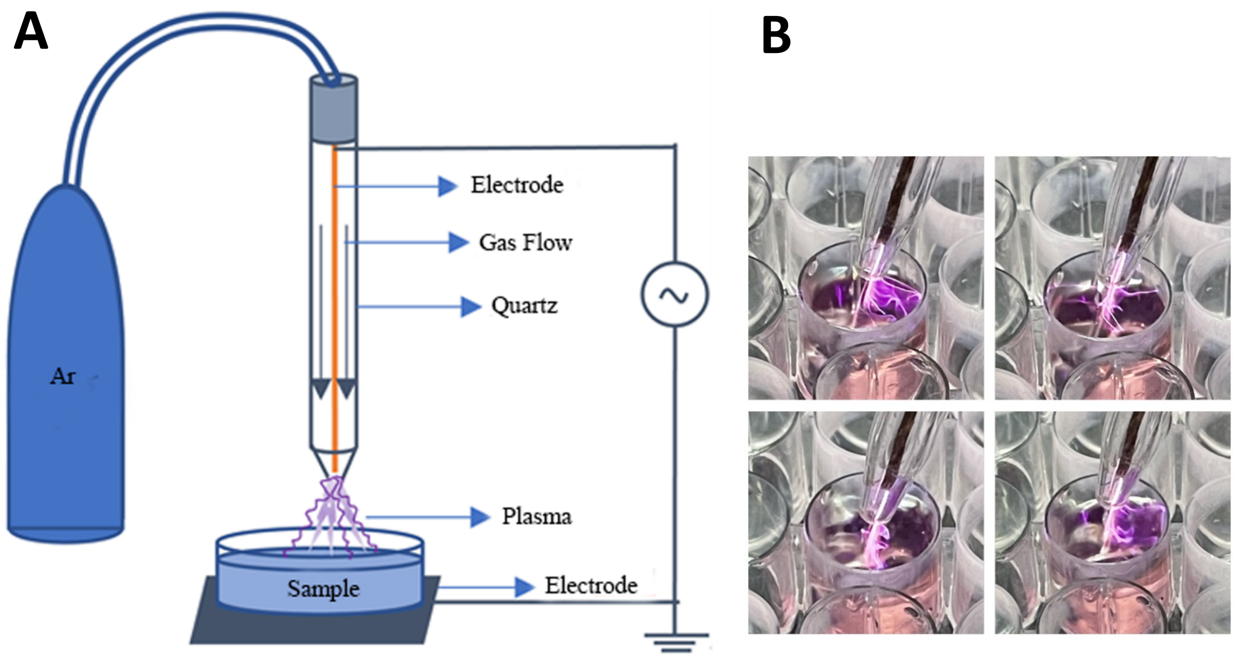


***Figure S2:*** *A) Schematic of the micro-discharge plasma setup consisting a 15 cm high quartz cylinder and a copper central electrode. An aluminum sheet was placed under the petri dish containing the sample as our ground electrode. B) Picture of plasma arc.*

We did the simultaneous synthesis and conjugation process using each of these setups, with three different gases, argon, helium, and oxygen, for three time periods of 5, 10, and 15 minutes and compared the results. Finally, according to the ELISA results showing the antigen and the antibody interaction, as well as considering the temperature changes, we chose the DBD setup with argon gas for 15 minutes. The results of the tests are given below.

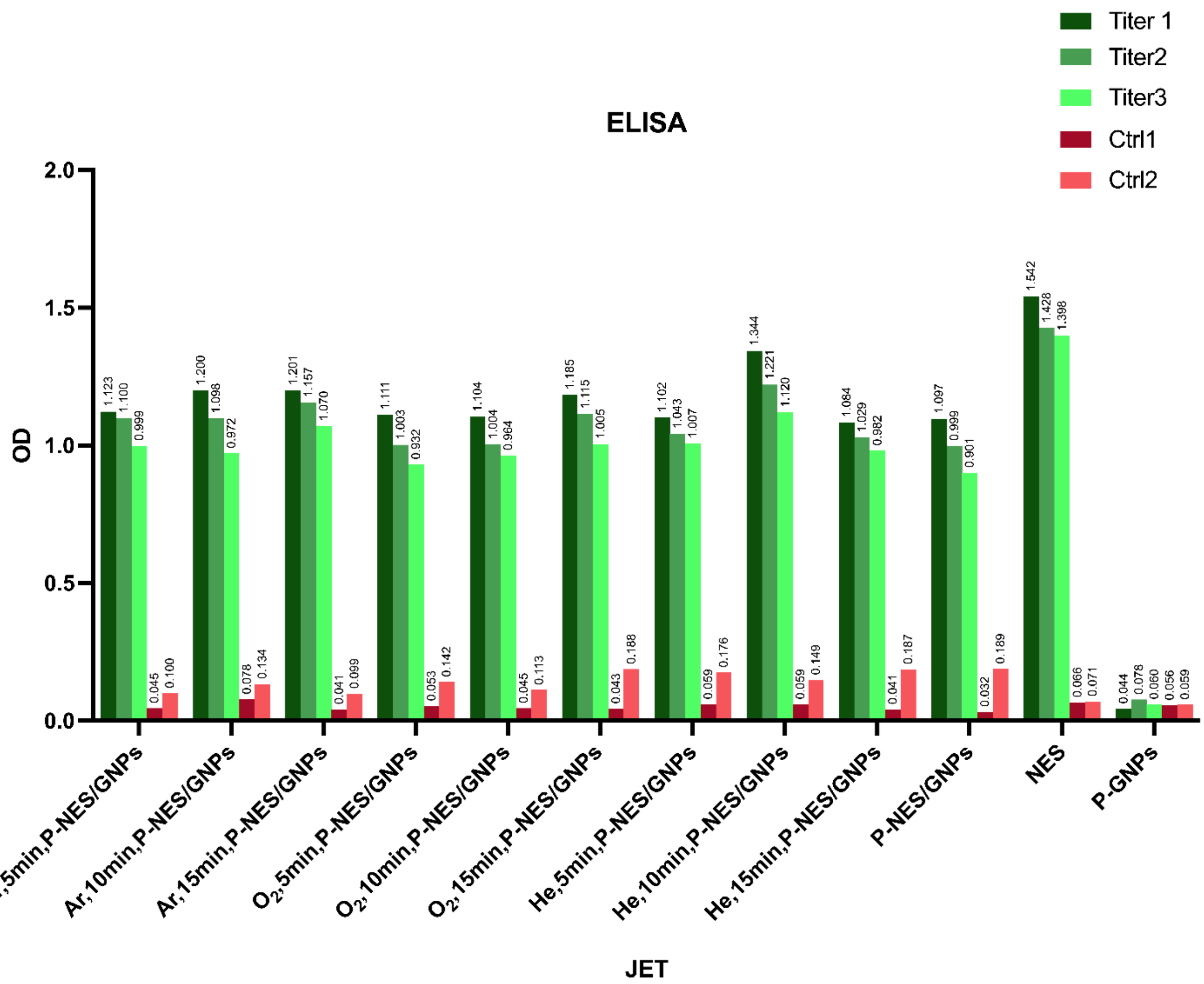


***Figure S3:*** *This chart presents the results of ELISA for the samples treated with plasma jet setup using three different gases: argon, oxygen, and helium. Samples were exposed to the plasma for three durations of 5, 10, and 15 minutes to assess the interaction between the antigen and antibody. This step was undertaken to determine the most effective gas and treatment time for plasma. For the jet plasma setup, we discovered that treating the samples with helium gas for 10 minutes yielded the most favorable outcomes.*

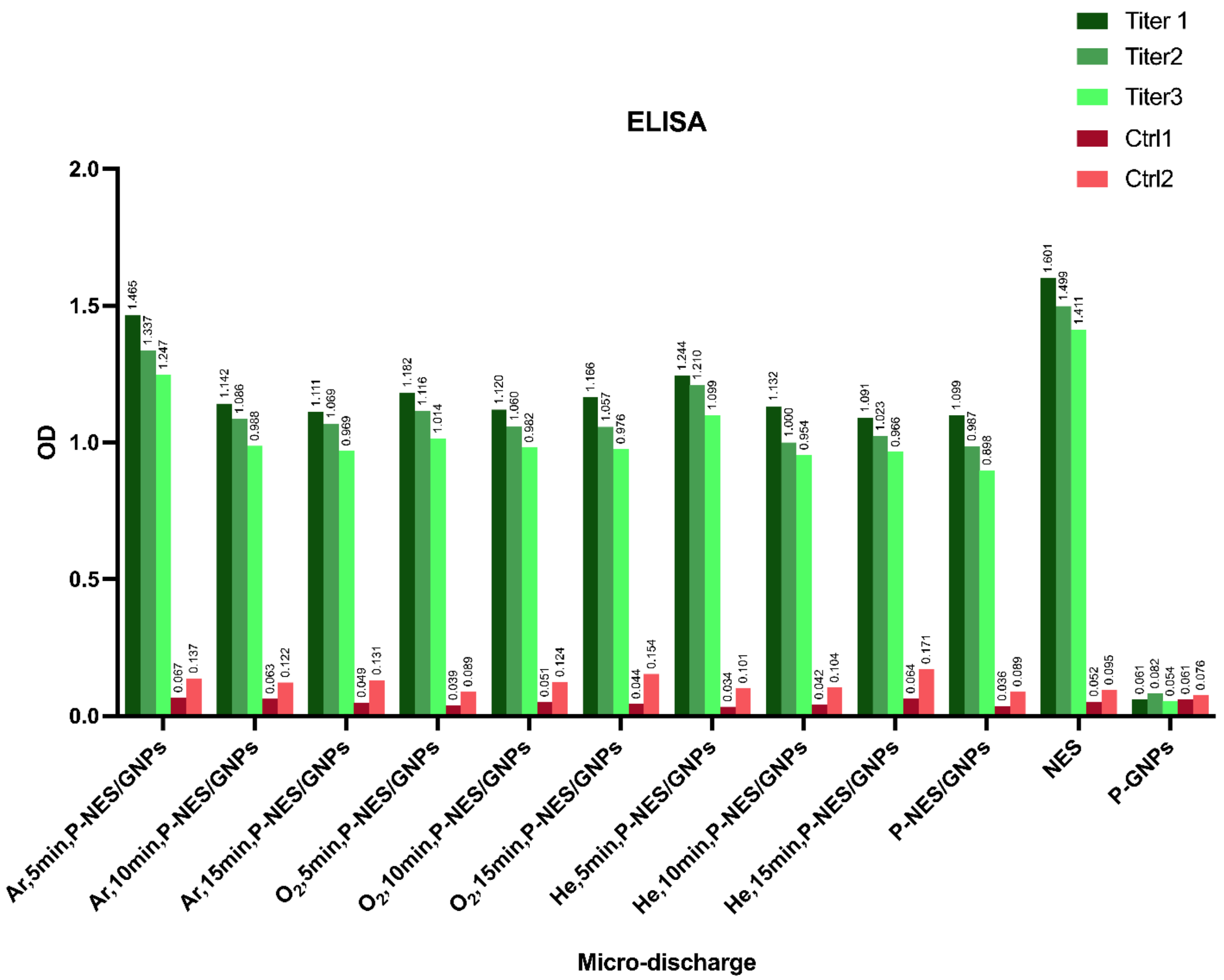


***Figure S4:*** *This chart compares the results of ELISA test on samples treated with micro-discharge plasma setup using three different gases (argon, oxygen, and helium) and three different treatment times (5, 10, and 15 minutes). The results indicate that the optimal combination of gas and treatment time for the micro-discharge plasma setup is argon gas for 5 minutes.*

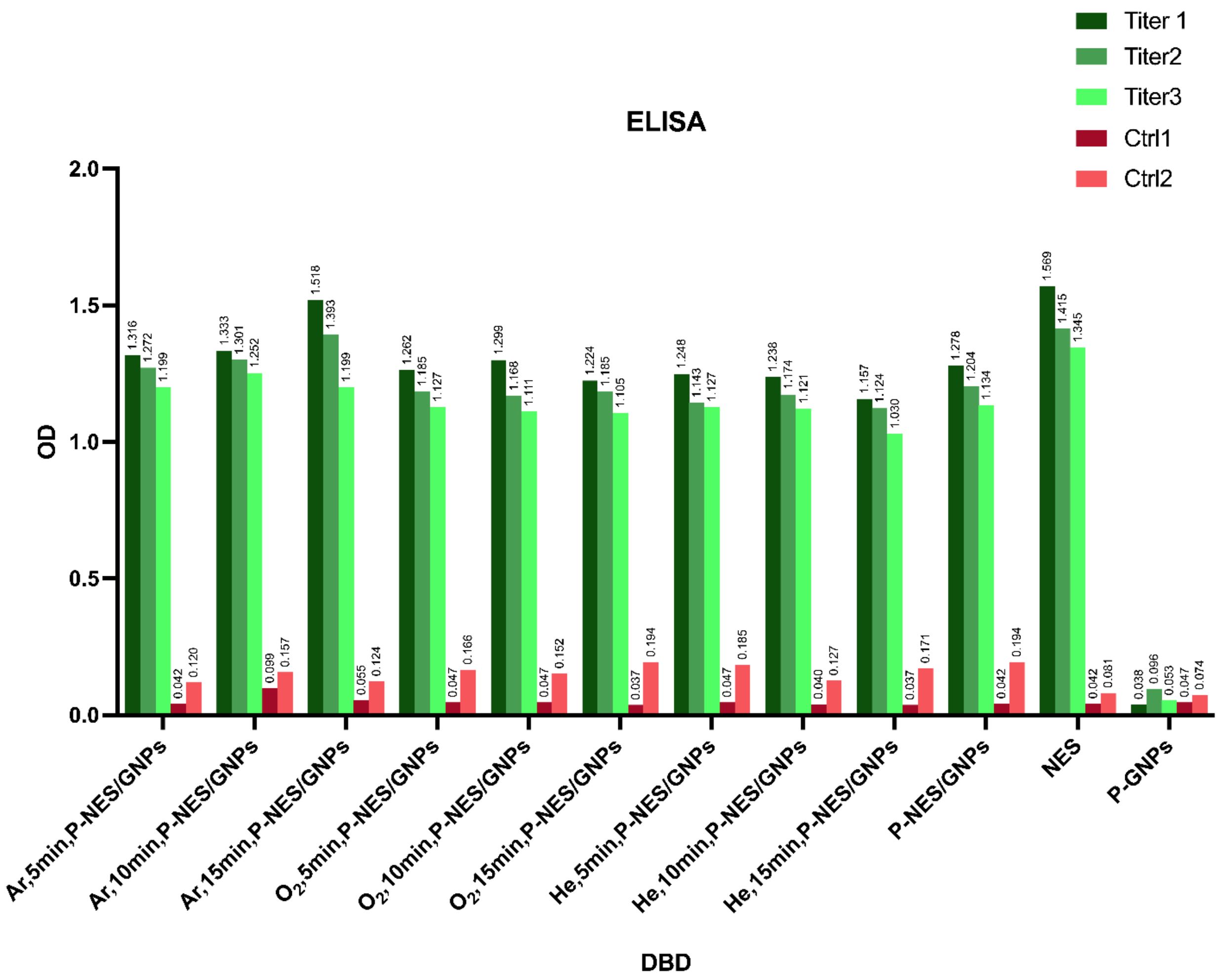


***Figure S5:*** *The chart evaluates the DBD plasma treatment's influence on ELISA assay outcomes, showing the antigen-antibody interaction when employing three distinct gases (Ar, $O_2$ and He) and varying treatment durations (5, 10, and 15 minutes). The data demonstrate that Argon gas treatment for 15 minutes elicited the most optimal response in the ELISA assay, suggesting this combination as the preferred parameter for the DBD plasma setup within this experimental context.*

The primary ELISAs presented here (Fig. 3, 4 and 5) were taken by indirect method, the details of which will be elaborated on later. To ensure result consistency, these ELISAs were repeated using the direct method as well. Since there was no difference in our results, with a large number of samples requiring analysis and the speed advantage of the direct method, the main testing phase transitioned to this approach. The direct ELISA protocol will be described in subsequent sections.

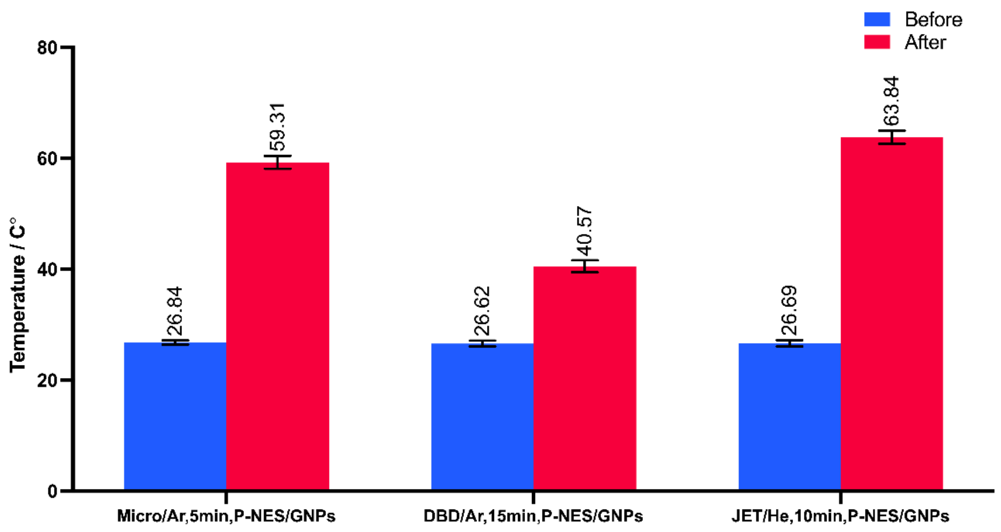


***Figure S6:*** *Considering that high temperature may affect the structure of proteins (including antigens) and can cause denaturation of antigens, temperature control was a crucial factor in our selection of the most suitable plasma setup. We sought a plasma setup that would induce the least temperature rise in our samples. To this end, we have investigated the temperature changes of the favorable samples from the previous parts. We repeated the temperature measurement of the samples three times. The results indicated that the DBD plasma setup exhibited the lowest temperature increase among the evaluated conditions. Consequently, considering the overall findings, we opted to utilize the DBD setup for our main test, employing argon gas for a treatment duration of 15 minutes.*

**Section 2:**

**Filtration**

To demonstrate the successful conjugation of the NES with GNPs, it was essential to eliminate any unbound, unconjugated antigens from our prepared samples. This was crucial to ensure that the antibody-antigen interaction observed in the ELISA assay was exclusively attributable to the conjugated antigen-GNP conjugates. To optimize the filtration step and achieve a balance between maximizing the removal of free antigens and minimizing the risk of aggregating or sedimenting the GNPs, a preliminary experiment was conducted to evaluate the effects of different filtration times.

In this experiment, naïve NES samples were filtered for 10, 20, and 30 minutes using a centrifuge at 1000 RPM. Following filtration, ELISA assays were performed on the filtrates (downstream and upstream of the filter) samples. The ELISA results for the 20- and 30-minute filtration GNPs included groups showed no significant difference, indicating that a filtration time of 20 minutes was sufficient to effectively remove free antigens without compromising the integrity of the GNPs conjugates (The ELISA test showed a slightly weaker optical density (OD) for the upstream of the filter in the 30-minute group compared to the 20-minute group, so we assumed that this time may cause GNPs to sedimenting). Therefore, a filtration time of 20 minutes was adopted for all main test samples to ensure the accurate assessment of the antibody-antigen interaction solely mediated by the conjugated antigens.

This optimization of the filtration step not only enhanced the accuracy of the ELISA assay but also provided valuable insights into the stability of the GNP conjugates. The ability to maintain the conjugation efficiency over prolonged filtration times suggests the robustness and stability of the conjugates, which is essential for their practical applications in targeted diagnostic and therapeutic strategies.

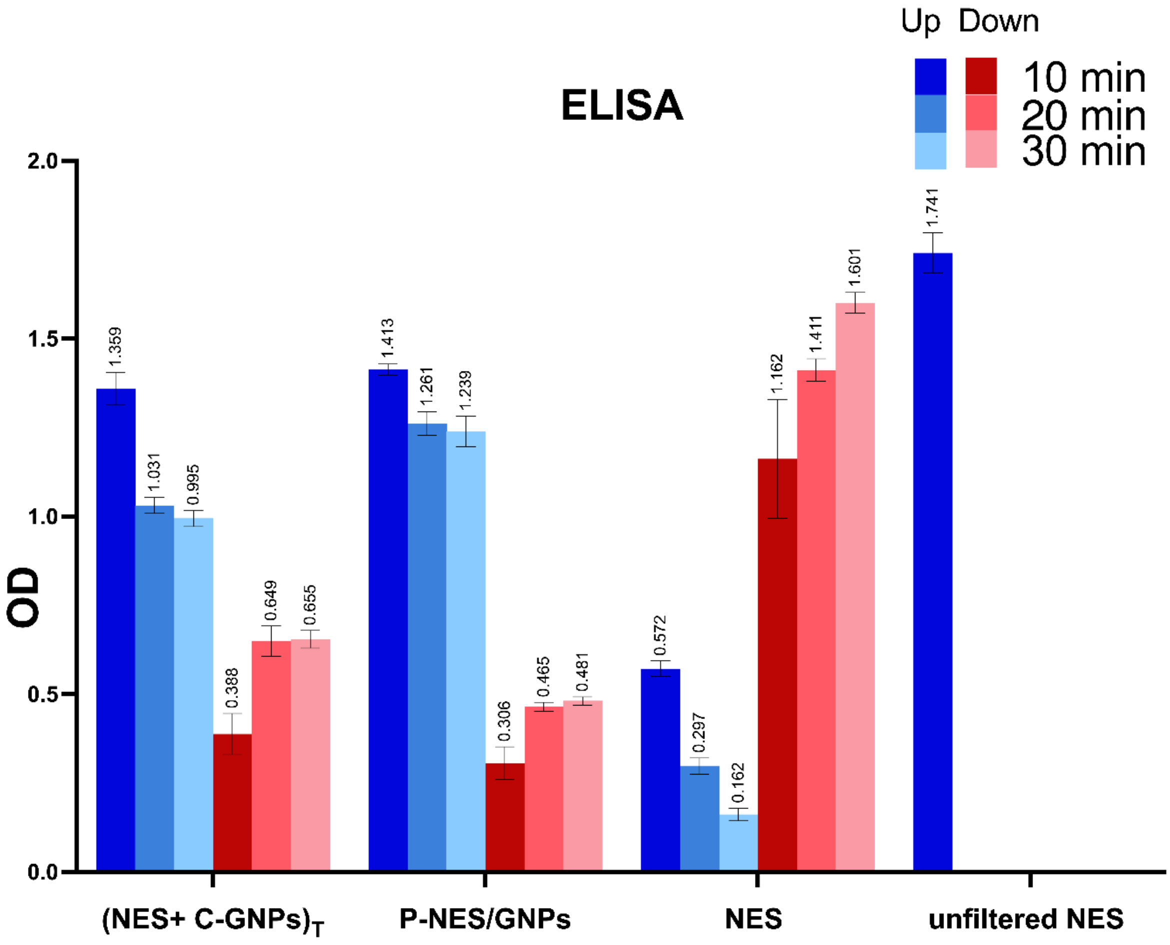


***Figure S7:*** *ELISA results of filtrates from 10-, 20- and 30-minute filtration groups in 3 times, revealing antigen removal efficiency while maintaining GNPs conjugate integrity. A filtration time of 20 minutes is deemed sufficient for effective filtration, minimizing sedimentation of GNPs. This chart summarizes the key findings of the ELISA results.*

**Section 3:**

**ELISA protocol**

Indirect ELISA Protocol:

1. Preparation:
    i. We coated 100 µL of our sample containing antigens onto ELISA wells.
    ii. Incubated the wells for 2 hours at 37°C.
2. Blocking and Washing:
    i. In this step we blocked unbound sites on wells with 300 µL 3% BSA solution and incubated the wells for 1.5 hours at 37°C after adding BSA.
    ii. Here we washed wells twice with PBS-Tween solution.
3. Primary Antibody Addition and Washing
    i. We added 100 µL of primary antibody to each well.
    ii. Then incubated it for 1 hour at 37°C.
    iii. Next, we washed wells three times with PBS-Tween solution.
4. Secondary Antibody Addition and Washing:
    i. We added 100 µL of secondary antibody (HRP-conjugated) per well.
    ii. Incubated the wells for 30 minutes at 37°C.
    iii. And washed wells four times with PBS-Tween solution.
5. TMB Addition and Reaction:
    i. We added 100 µL of TMB to each well.
    ii. We let the reaction proceed for 10 minutes.
    iii. Then stopped the reaction with 50 µL of 2M HCl as soon as control groups started to change color.
6. Finally, we read the OD of per well using an ELISA Reader at 450 nm – 630 nm.

Direct ELISA Protocol:

1. Preparation:
    i. We coated 100 μL of our sample containing antigens onto ELISA wells.
    ii. Incubated the wells for 2 hours at 37°C.
2. Blocking and Washing:
    i. In this step we blocked unbound sites on wells with 300 μL 3% BSA solution and incubated the wells for 1.5 hours at 37°C after adding BSA.
    ii. Then we washed wells twice with PBS-Tween solution.
3. Primary HRP-conjugated Antibody Addition and Washing
    i. We added 100 μL of primary antibody to each well.
    ii. Incubated the wells for 1 hour at 37°C.
    iii. Afterwards, we washed wells three times with PBS-Tween solution.
4. TMB Addition and Reaction:
    i. We added 100 μL of TMB to each well.
    ii. We let the reaction proceed for 10 minutes.
    iii. Then stopped the reaction with 50 μL of 2 M HCl as soon as control groups started to change color.
5. Finally, we read the OD of per well using an ELISA Reader at 450 nm – 630 nm.

**Section 4:**

**Stability**

Eventually, to assess the stability of the P-NES/GNPs and NES+(C-GNPs) samples, we subjected them to repeated UV-vis and ELISA analyses after one week (the time when the first ELISA was taken from our main samples), two weeks, and one month. The samples were stored at 4 degrees in the refrigerator. Our findings revealed minimal variations in the results obtained during these time intervals. The UV-vis spectra exhibited a slight decrease in absorbance after one month, potentially attributed to the deposition of a small quantity of nanoparticles. The ELISA results, which reflect the interaction of conjugated antigen with antibody, showed no significant difference in the OD readings throughout the study period. Based on these observations, it can be concluded that the simultaneous synthesis and conjugation (P-NES/GNPs) and linker-free plasma conjugation (NES+(C-GNPs)) methods yielded stable nanomaterial formulations, rendering them suitable for practical applications.

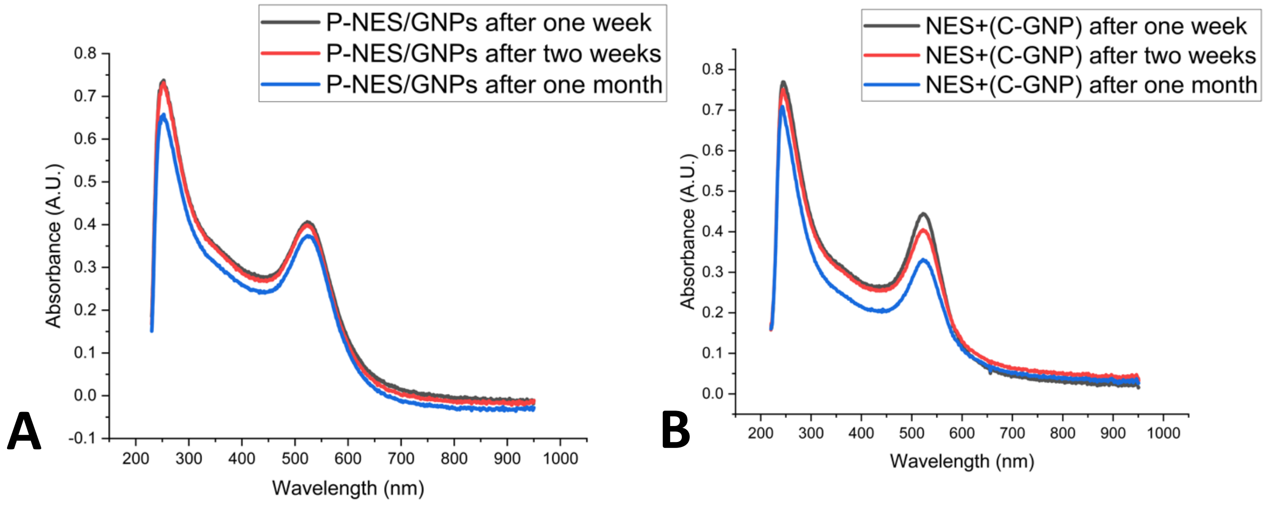


***Figure S8:*** *A) UV-vis taken from P-NES/GNPs. B) UV-vis taken from NES+(C-GNPs). Both showing the stability of samples after one week, two weeks, and a month.*

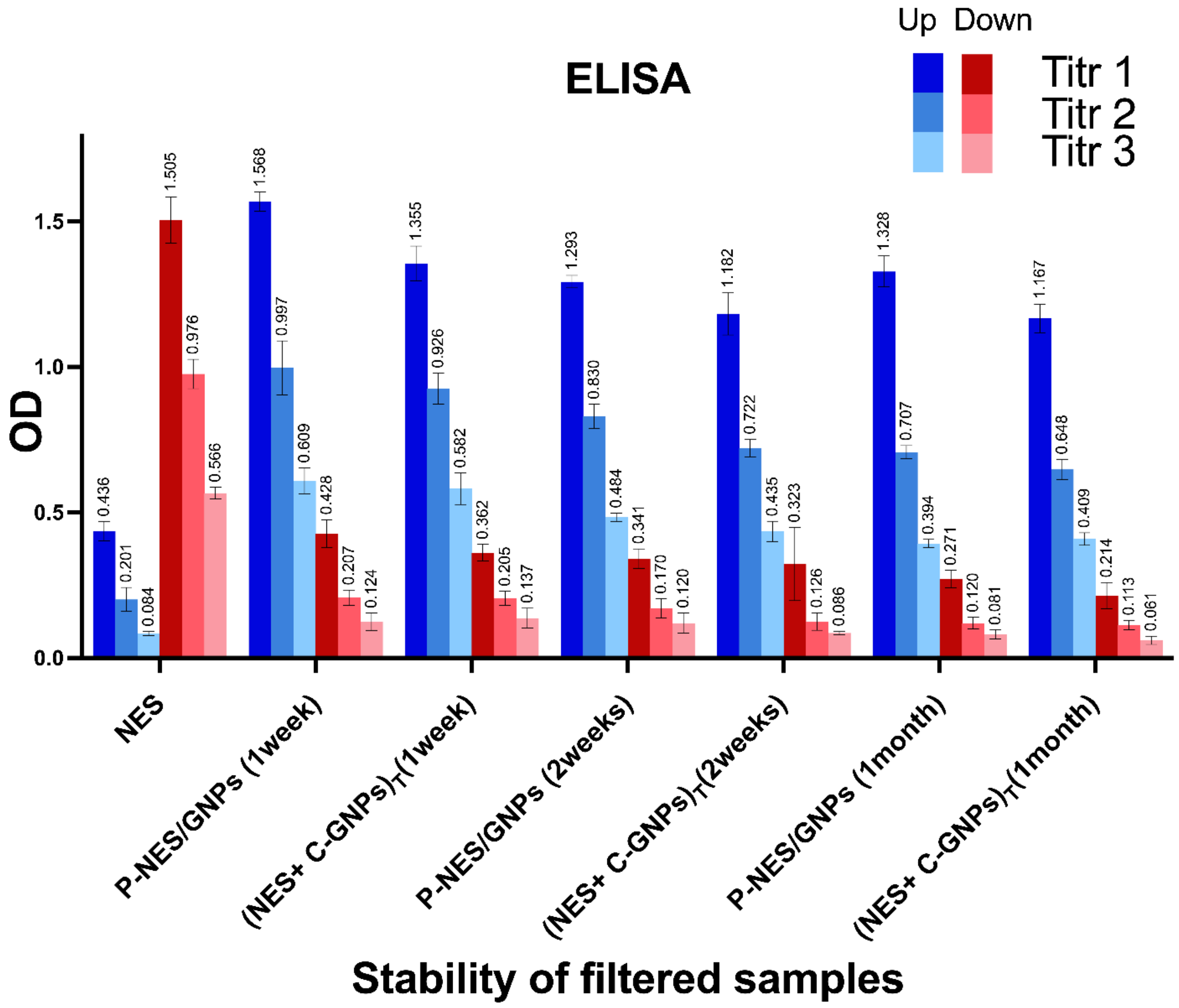


***Figure S9:*** *ELISA taken from P-NES/GNPs and NES+(C-GNPs) samples which indicates the stability of our conjugates after one week (first test), two weeks, and a month.*

**Section 5:**

**Efficiency**

To quantify the conjugation efficiency, we employed the following equation based on Figure 3.C (NES concentration graph) in the main text. All concentrations are calculated ± Standard Deviation (SD):

Filtered NES-Upstream: 5.35 ± 0.95 μg/ml

Filtered NES-Downstream: 40.97 ± 1.97 μg/ml

Filtered P-NES/GNPs-Upstream: 43.08 ± 0.79 μg/ml

Filtered P-NES/GNPs-Downstream: 4.84 ± 1.18 μg/ml

$$\text{Efficiency} = \frac{\text{Filtered P.NES/GNPs.UP(conc)} - \text{Filtered NES.Up(conc)}}{\text{Unfiltered NES(conc)} - \text{Filtered NES.Up(conc)}} * 100 =$$

$$\frac{(43.08 \pm 0.79) - (5.35 \pm 0.95)}{50 - (5.35 \pm 0.95)} * 100 = 84.55 \pm 6.65\ \%$$

The amount of naïve NES in 1 ml that did not pass through the filter after the optimum time is subtracted from the amount of NES conjugated to GNPs at the upstream of the filter in 1 ml. In the denominator of the fraction, the total amount of NES (50 μg) in 1 ml minus the amount of naïve NES in 1 ml, which did not pass through the filter (considered as the background of our test), is included. Thereby, we have calculated the efficiency of our work.

**Section 6:**

**Zeta-sizer**

The size of nanoparticles plays a crucial role in their biological interactions and therapeutic efficacy. Dynamic light scattering (DLS) and transmission electron microscopy (TEM) are widely used techniques to determine the size of nanoparticles. However, the results obtained from these techniques may diverge due to the distinct physical principles they employ.

According to Einstein's Stokes equation, the diffusion coefficient, a measure of the particle's movement due to Brownian motion, is inversely proportional to the hydrodynamic radius. This means that smaller particles diffuse more rapidly than larger particles.

In DLS, the scattered light from nanoparticles undergoes a frequency shift due to the interaction between the light and the particles. This frequency shift is directly related to the particle's diffusion coefficient and, consequently, its hydrodynamic radius. Therefore, a smaller hydrodynamic radius translates to a larger frequency shift.

TEM provides direct visualization of the nanoparticles, revealing their physical dimensions. However, DLS measures the hydrodynamic radius, which includes the effects of the surrounding medium. The adsorbed water molecules around nanoparticles can alter their apparent size, leading to a larger hydrodynamic radius compared to the physical radius observed in TEM.

The placement of an antigen chain on the surface of nanoparticles can further influence their hydrodynamic properties. The presence of the antigen chain can induce the adsorption of additional water molecules around the nanoparticles, significantly increasing their hydrodynamic radius. This effect is particularly evident for P-NES/GNPs compared to P-GNPs. TEM revealed an average size of 75±15 nm for P-GNPs, while DLS indicated an average size of 80±10 nm. This difference is more pronounced for P-NES/GNPs, with TEM suggesting an average size of 20±5 nm and DLS measuring an average size of 29±10 nm. The NES presence has enhanced the hydrophilicity of the GNPs, leading to increased water adsorption and a more dispersed particle size distribution as observed by DLS.

The hydrodynamic radius of nanoparticles plays a crucial role in their interactions with biological systems. Larger hydrodynamic particles tend to exhibit enhanced circulation and accumulation in specific tissues, potentially improving their therapeutic efficacy. However, the increased hydrodynamic size can also lead to aggregation and reduced cellular uptake, compromising their therapeutic potential.

In summary, the hydrodynamic radius, diffusion rate, and frequency shift are interconnected parameters that provide valuable insights into the size and behavior of nanoparticles. The discrepancy between TEM and DLS results emphasizes the importance of considering both physical and hydrodynamic size in nanoparticle characterization.

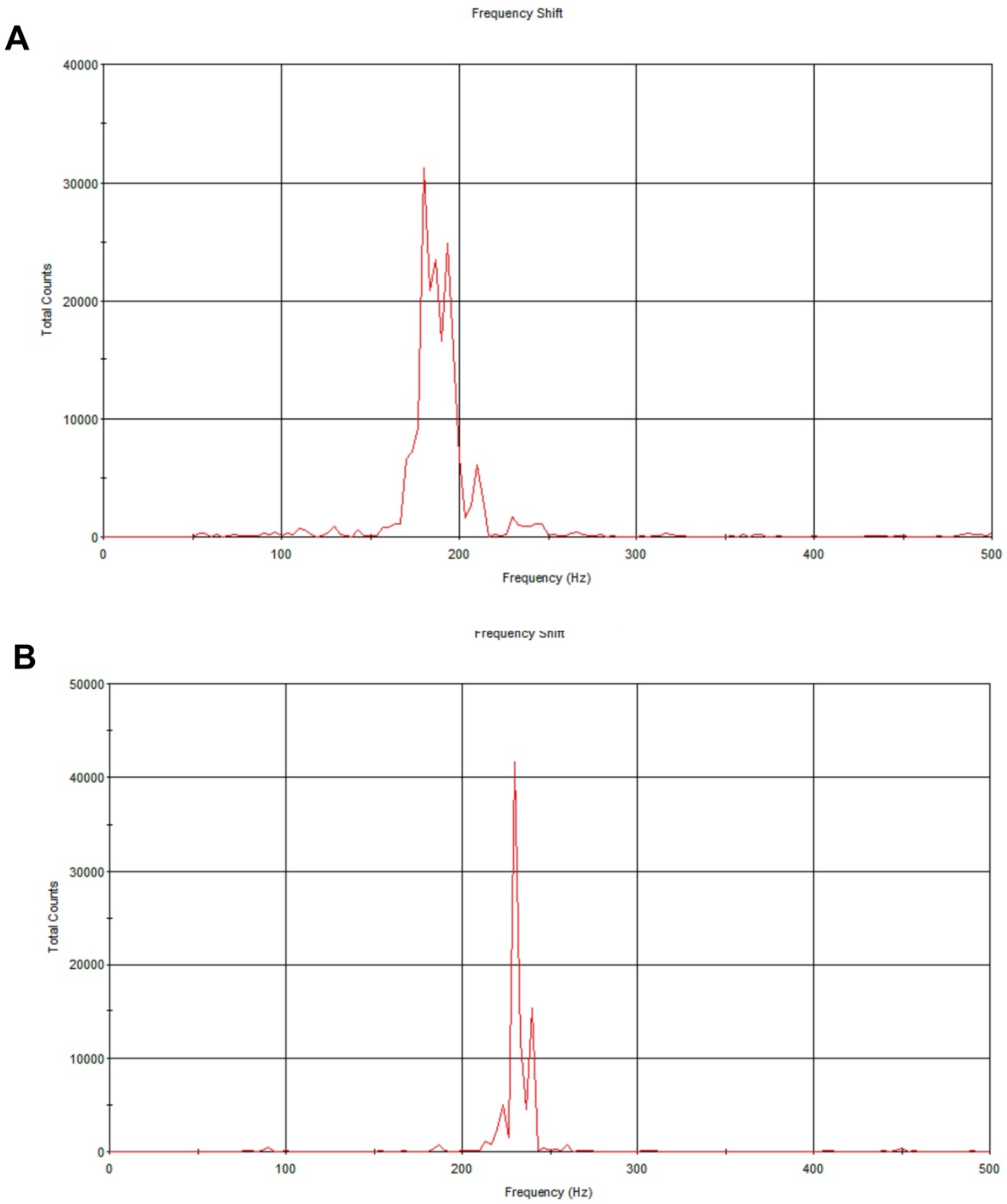


***Figure S10:*** *A) Demonstrating the frequency shift distributions for P-GNPs B) The frequency shift distributions for P/NES-GNPs. The P/NES-GNPs group exhibits a higher peak frequency shift, demonstrating the enhanced particle mobility upon antigen conjugation*

.

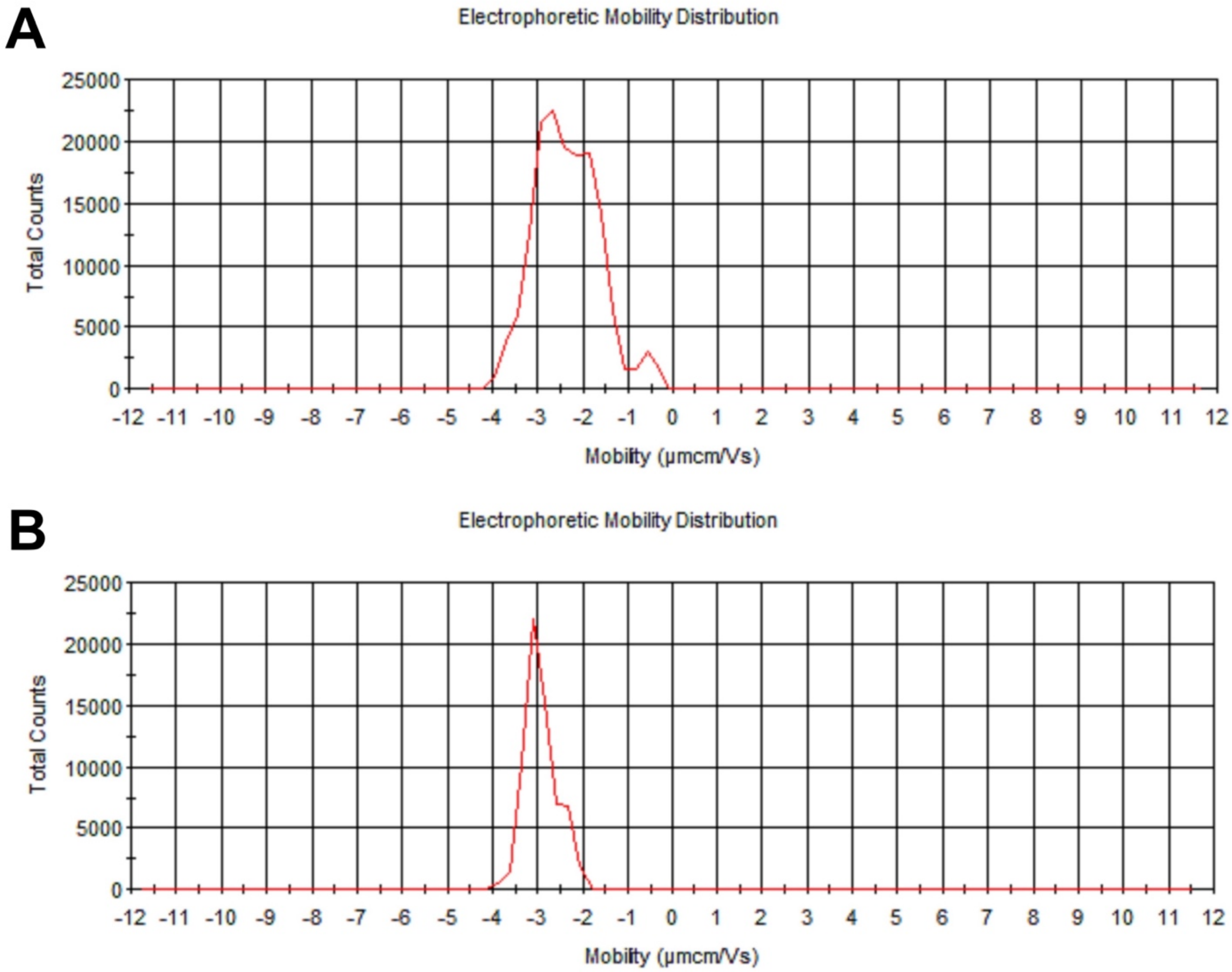


***Figure S11:*** *A) Demonstrating the Electrophoretic mobility of P-GNPs B) P/NES-GNPs Electrophoretic mobility. As it is obvious the graph shows enhanced particle mobility upon antigen conjugation.*

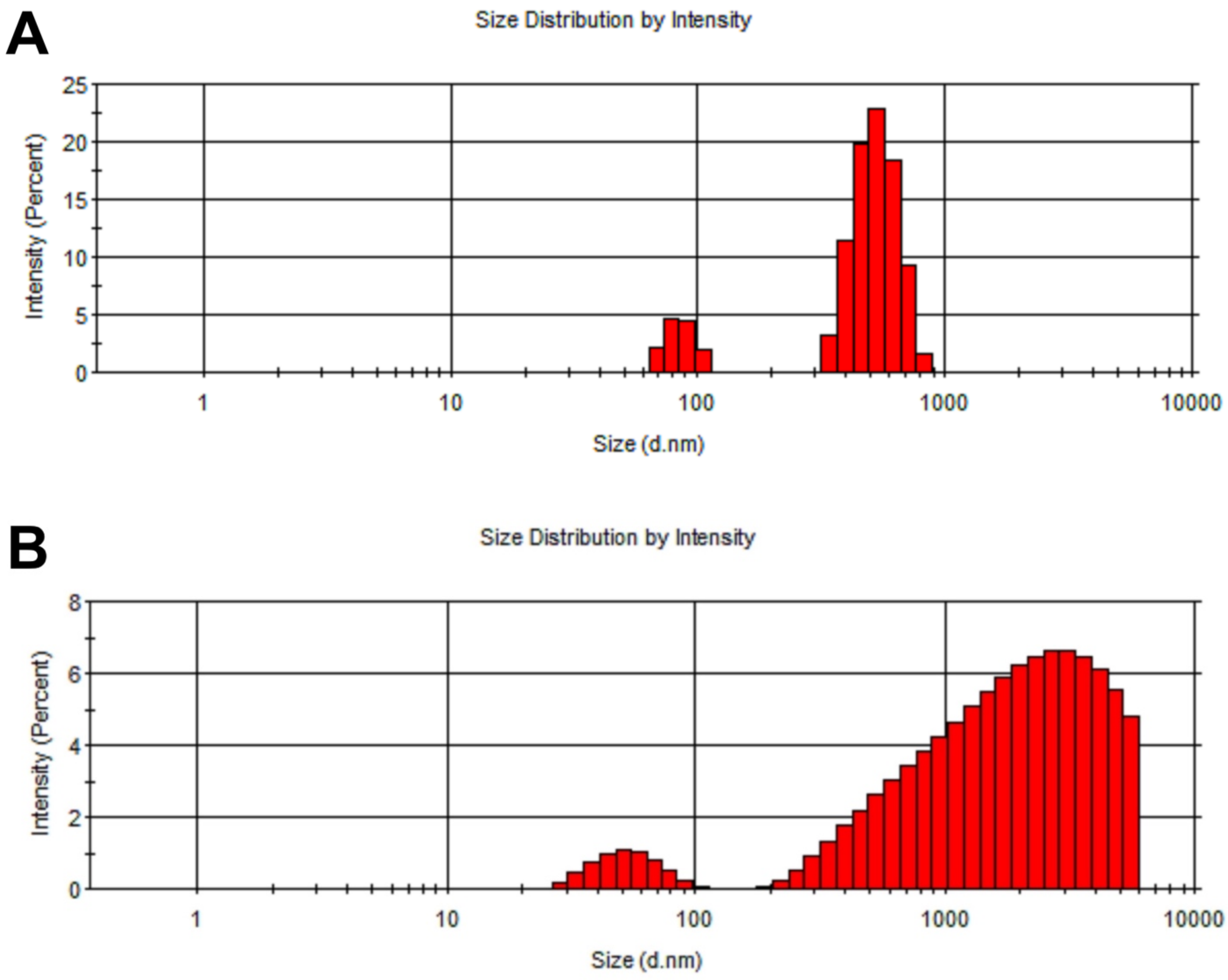


***Figure S12:*** *These graphs show the particle size distribution based on the intensity for A) P-GNPs and B) P-NES/GNPs. According to Figures A and B, low peaks of approximately 30 nm and 85 nm are observed, respectively, along with a relatively wide and high peak at dimensions above 1000 nm. As no peak was observed in the size distribution diagram in terms of the number in this size range, it can be concluded that a very small number of particles may have aggregated. The peaks around 85 nm for GNPs and around 30 nm for P-NES/GNPs confirm our UV-vis and TEM results regarding the particle sizes.*

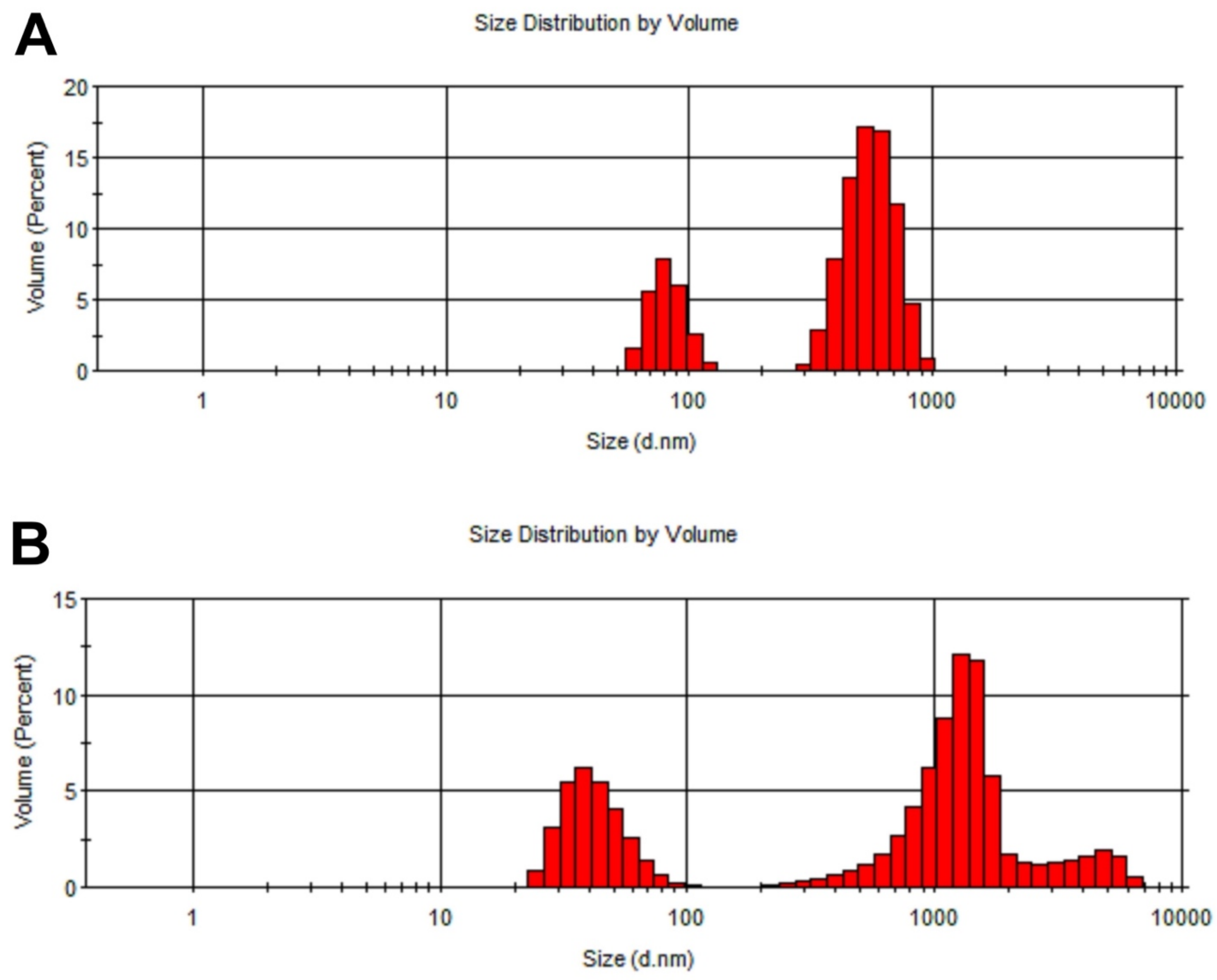


***Figure S13:*** *These graphs show size distribution based on volume. A) P-GNPs and B) P-NES/GNPs. With a similar explanation for the size distribution by intensity diagram, a peak at about 80 nm is observed for P-GNPs. Additionally, for P-NES/GNPs, a peak is visible at less than 40 nm. The difference in size compared to what was observed in TEM is attributed, as explained, to the influence of the hydrodynamic radius in the DLS test. Moreover, the presence of a peak around 700 to 1100 nm indicates a small number of aggregated particles (as there is no peak in this area in the size distribution by number graph provided in the main text).*
